# Evaluation of rubber tree transcriptome and discovery of SNP and SSR from candidate genes involved in cellulose and lignin biosynthesis

**DOI:** 10.1101/2021.09.02.458661

**Authors:** Ahmad Sofiman Othman, Mohd Fahmi Abu Bakar

## Abstract

*Hevea brasiliensis* (the rubber tree) is a well-known species with high economic value, and it is the primary source of natural rubber globally. Increasing demand for furniture and related industries has made rubberwood production as important as latex production. Molecular markers such as Single Nucleotide Polymorphisms (SNPs) and Simple Sequence Repeats (SSRs) are widely used for Marker Assisted Selection (MAS) which can be detected in large quantity by transcriptome sequencing. MAS is thought to be a useful method for the development of new rubberwood clones for its shorter breeding cycle compared to a conventional breeding procedure. In this study we performed RNA sequencing (RNA-seq) on four *H. brasiliensis* clones (RRIM 712, RRIM 2025, RRIM 3001 and PB 314) from three tissues including bark, latex and leaf samples to identify SSRs and SNPs associated with wood-formation related genes. The RNA sequencing using the Illumina NextSeq 500 v2 platform, generated 1,697,491,922 raw reads. A total of 101,269 transcripts over 400 bp in size were obtained and similarity search of the non-redundant (nr) protein database returned 83,748 (83%) positive BLASTx hits. The transcriptome analysis was annotated using the NCBI NR (National Center for Biotechnology Information Non-Redundant), UniProtKB/Swiss-Prot, gene ontology (GO), and Kyoto Encyclopedia of Genes and Genomes (KEGG) databases. Differential expression analysis between later-timber rubber clone and non-later-timber rubber clone on wood-formation related genes, showed genes encoding phenylalanine ammonia-lyase (PAL), caffeic acid O-methyltransferase (COMT) and cinnamoyl-CoA reductase (CCR) were highly up-regulated in a latex-timber rubber clone. In total, about 3,210,629 SNPs and 14,956 SSRs were detected with 1,786 SNPs and 31 SSRs were found for wood-formation biosynthesis of H. brasilensis from 11 lignin and cellulose gene toolboxes. After filtering and primer selection, 103 SNPs and 18 SSR markers were successfully amplified and could be useful as molecular tool for marker assisted breeding to produce new timber rubber clones.

## Introduction

*Hevea brasiliensis* (Wild.) Muell-Arg. is a tree native to the Brazilian Amazon region and from the Euphorbiaceae family [1]. This species is the most abundant species of the genus, with the largest production capacity, accounting for about 99% of all natural rubber produced globally and with the greatest genetic variability [2]. In the past, a greater emphasis was put on production of only high-latex-yielding clones, giving rise to a spectacular increase in yield. However, the increasing demand for rubberwood furniture has given rise to a new identity for the rubber industry [3]. Rubberwood furniture (advertised as ‘environmentally friendly’) [4], which can be produced from latex-timber clones, requires considerable growth time using conventional breeding [5]. Due to its relatively low cost and light colour, rubberwood furniture has gained wide acceptance amongst domestic and foreign consumers and is now accepted as being a viable alternative to other timber species [6].

Recently, the RRIM 3001 clone has been successfully bred with a girth size almost double that of other clones, with increased wood production [5]. In wood formation, lignin and cellulose are the main components of plants’ secondary wall [7]. Several genes are responsible for the biosynthesis of lignin and cellulose. However, there is less studies focussing on the lignin and cellulose biosynthetic genes at genomic levels. The lack of information regarding the expression of genes related to lignin and cellulose biosynthesis can influence the development of new bred with high wood production characteristics. Therefore, genetic information is very important to be gathered via sequencing *Hevea* genomic data, and these data are essential for obtaining information at the chromosome/genome level and for discovering novel scientific knowledge about rubberwood traits. Genomic data can also improve the quality of breeding programmes by enhancing agronomic traits of commercial importance.

RNA-Seq could potentially be a useful tool for understanding the biological process in *Hevea* governing agronomic traits, including rubberwood quality. High-throughput genomic techniques associated with innovative bioinformatics tools can help in rubber-tree breeding and can facilitate the development of superior clones [1]. With the reduced cost of next-generation sequencing (NGS) technologies, RNA sequencing (RNA-seq) has become widespread because it enables high-resolution characterization of transcriptomes. This method has many advantages, including single-base resolution, enabling detection of thousands of single-nucleotide polymorphisms (SNPs) and simple-sequence repeats (SSRs) for further development of SNP and SSR markers [8]. These markers can be useful for functional saturation of linkage maps and can also be used for marker-assisted selection (MAS).

In this study, we utilized paired-end Illumina sequencing on selected rubber clones to identify the genes involved in cellulose and lignin biosynthesis in this non-model tree. Using a standard mapping algorithm, we examined the quality of contigs generated and tried to identify cellulose- and lignin-biosynthesis genes. Our main purpose was to detect SNPs and SSRs within genes controlling wood-related traits. As our study involved multiple rubber clones (namely RRIM 712, RRIM 2025, RRIM 3001 and PB 314) with differing rubberwood quality, it could provide a useful tool for the selection of offspring with favourable traits in breeding programmes.

## Materials and methods

### Plant materials and total RNA isolation

Bark, leaf, and latex tissues were obtained from a 10-year-old rubber tree (*H. brasiliensis* clone; RRIM 712, RRIM 2025, RRIM 3001 and PB 314), located at (Rubber Industry Smallholders Development Authority) RISDA Semaian, Bukit Perak, Pendang, Kedah, Malaysia. RRIM 712 is a latex clone, while RRIM 2025, RRIM 3001 and PB 314 are latex-timber clones. The girth size of each clone was different: 42-44 cm in the case of RRIM 712; 50-52 cm in the case of RRIM 3001; 45-47 cm in the case of PB 314; and 47-49 cm in the case of RRIM 2025. The collected samples were frozen in liquid nitrogen and transferred to the laboratory in dry ice. The total RNA from leaf, bark and latex was extracted following the Qiagen RNAeasy Plant MiniKit protocol (Qiagen Inc., Chatsworth, CA). RNA quality and integrity were determined using an Agilent 2100 Bioanalyzer (Agilent Technologies, Palo Alto, CA).

### cDNA library construction and sequencing

The high quality of extracted RNAs with OD A260/A230 value of more than 2.0, OD A260/A280 value of between 1.8 to 2.0 and concentration of more than 100ng/uL were used to construct paired-end Illumina mRNA libraries using the Illumina NextSeq 500 v2 Kit. Each sample was sequenced in multiple HiSeq2000 lanes using the NextSeq 500 v2 Cycle Kit (Illumina, San Diego, CA) in order to obtain 2 × 75-bp reads.

### Raw data filtering

Raw reads generated in FASTQ format (obtained from Illumina platforms) were analysed using FastQC software [9] resulting in clean reads which will be used for mapping and assembly. Clean reads were obtained by filtering the raw reads based on these steps that include: 1) removing reads containing adaptors; 2) removing reads containing ambiguous nucleotides; 3) trimmed low-quality reads to exclude poor-quality reads. FastQC produces end results as clean reads, and these were to be used for further analysis.

### Transcriptome mapping and assembly

Once clean reads were obtained, they were mapped onto the draft *H. brasiliensis* genome [10] (accession: PRJDB4387) by Bowtie2, via the DDBJ/EMBL/GenBank BioProject database, using TopHat (version 2.1.0) software [11]. The reads were mapped onto the genome with default parameters. The mapped reads were then assembled using Cufflink v2.2.1, with default parameters. To obtain isoforms with a high degree of confidence, only those assembled transcripts with a value of FPKM > 0 were retained for further analyses. The assembled transcript sequence generated by Cufflink from the rubber-genome sequence was then extracted. Unmapped reads were extracted from the unmapped BAM file. *De novo* assembly of the reference reads was performed using Trinity v2.6.5 [12] with default parameters in order to generate the reference transcriptome. Unmapped reads were then mapped to the reference transcriptome using TopHat (version 2.1.0) software [11]. The assembled sequences generated by Cufflink from genome mapping and reference-transcriptome mapping using TopHat were combined. Finally, Cd-hit software was used [13] to combine assembled sequences from both sources and to reduce redundancy of the sequences. The output transcripts were considered as *Hevea* reference sequences.

### Gene annotations

Basic annotations were executed that include protein functional annotation following searches of several databases, namely the NCBI NR (National Center for Biotechnology Information Non-Redundant) database, UniProtKB/Swiss-Prot (Universal Protein Resources) database and KEGG (Kyoto Encyclopedia of Genes and Genomes) database. Subsequently we performed GO (Gene Ontology) functional annotation of unigene using Blast2GO software [14], set up with an e-value cut-off at 1^e-10^. The functional classification of unigene was done by WEGO (version 2.0) software [15].

### SSR markers detection and primer designing

We identified total SSRs and SSRs from lignin and cellulose genes toolboxes within each rubber genotypes using the microsatellite prediction (MISA) program (http://pgrc.ipk-gatersleben.de/misa/). Once SSRs were identified, the corresponding primers were designed by using the following parameters: (1) motif ranged from dinucleotides to pentanucleotides (2) minimum repeat unit was five for all nucleotide motifs (3) the maximum interruption lengths between two SSR markers was set up as 100bp. We used the program Primer3 (http:/primer3.ut.ee/) to design primers using the criteria of GC contents for primer sequences ranging from 40% to 60%.

### SNP identification and primer designing

For SNP detection, we used SAMtools v1.3.1 [16] first to generate mpileup for one or multiple BAM files. Subsequently VarScan v2.3.9 [17] was then used to perform total SNPs and SNPs from lignin and cellulose genes toolboxes detection using default parameters. Reads were mapped against the reference transcriptome and filtered by ascertaining which DNA bases were different from the reference. Selected SNPs were required to have a read depth equal to or greater than 10, an SNP reads/total reads ratio equal to or greater than 0.25, a minimum Phred score of 20 and an SNP quality of 50. Similarly, we used Primer3 (http:/primer3.ut.ee/) to design primers as mentioned in section above.

### Molecular-marker amplification and sequencing

The SSRs and SNPs loci associated with lignin and cellulose gene toolboxes were amplified. Amplified SSRs were sequenced to ascertain their workability. PCR amplifications for SSRs and SNPs loci were performed in 25-ul reactions that includes buffer solution, dNTP, MgCl and *Taq* polymerase using Mytaq Master Mix (Bioline). The PCR reaction condition is as follows: initial denaturation at 95°C for 2 min, followed by 35 amplification cycles (30 s at 95°C, 30 s at the specific annealing temperature and 1 min at 72°C), and a final extension at 72°C for 10 min. The PCR products were resolved via electrophoresis in 1.5% agarose gels prior to identifying the specific amplification SSRs and SNPs loci. The DNA sequencing reaction was performed by the service provider First Base company. The sequencing data were then visually inspected using Molecular Evolutionary Genetics Analysis (MEGA) X software [18].

### Phylogenetic Analysis

The determination of genes relationships related to lignin biosynthesis between rubber clones was done by using phylogenetic analysis. Four genes involved in lignin biosynthesis; from leaf tissue of RRIM 712, RRIM 2025, RRIM 3001 and PB 314 including caffeic acid O-methyltransferase (COMT), caffeoyl-CoA O-methyltransferase (CCoAOMT), cinnamoyl-CoA reductase (CCR) and cinnamyl alcohol dehydrogenase (CAD) were amplified using Polymerase Chain Reaction (PCR) process with two replicate from each clones. The DNA sequence were then sent for sequencing. The sequence data were aligned using MEGA X software [18]. The aligned sequences were then used as an input to construct phylogenetic tree using Maximum Parsimony (MP) approach. MP approach was selected due to the criterion ability to weighted equally and unordered the character state changes where the changes occur as early in relation to the root as possible. The evaluation of confidence values of the clades was carried out through ten thousand replicates of the bootstrap analysis. The phylogenetic tree analysis was done using MEGA X software [18]. The phylogenies were rooted using outgroup species from *Mahihot esculenta* available on NCBI database.

## Results

### Transcriptome Analysis and Functional Annotation

In total, 1,697,491,922 raw reads were generated and trimmed to exclude low-quality reads. Of the 1,697,491,922 raw reads, 1,641,112,636 high-quality reads could be recovered after the trimming process. High-quality reads were then mapped onto the available draft *Hevea* genome (accession: PRJDB4387) and assembled to generate a total of 101,269 transcripts. The transcripts were considered as the reference transcriptome. The library of raw and clean reads is shown in S1 Table. The reference transcriptome lengths ranged from 424 to 10,503 bp. Of the 101,269 transcripts, 50,375 (49.7%) ranged in size from 1 to 2 kb, and 25,940 (25.6%) were longer than 2 kb. A summary of transcriptome assembly is shown in Table 1. All data are deposited under NCBI Genbank Bioproject available at https://www.ncbi.nlm.nih.gov/bioproject/PRJNA511923. The individual data identification is available under accession number between SRX5181842 to SRX5181865.

**Table 1.** Summary of the transcriptome assembly of four *H. brasiliensis* clones.

| Summary | Number of Transcripts |
| --- | --- |
| Number of transcripts | 101,269 |
| Mean transcript length | 1,046 |
| Maximum transcript length | 10,503 |
| Minimum transcript length | 424 |
| N50 length | 1,313 |

The 101,269 transcripts were searched against the UniProtKB/Swiss-Prot database using BLASTx, employing a cut-off e-value of 1e-10 as the criterion for defining a significant hit. Of these transcripts, 97,454 (96.2%) showed significant BLASTx matches in the UniProtKB/Swiss-Prot database. Moreover, the top five species with the highest number of hits in the NR database (annotated using BLASTx) included *Manihot esculenta, Jatropha curcas, Ricinus communis, Hevea brasiliensis* and *Populus trichocarpa* (S1 Fig). The 97,454 transcripts showing positive hits in the UniProtKB/Swiss-Prot database were then annotated using gene ontology (GO) terms. Of the 97,454 transcripts, 71,902 had hits in the GO database, while 25,552 transcripts were not assigned to any GO category (S2 Fig). Of the three main ontologies within the GO domain, biological processes were most highly represented (with 37,455 transcripts), followed by molecular functions (35,816 transcripts) and cellular components (34,155 transcripts). About 23 GO classification categories were identified for biological processes where metabolic processes (38,701 transcripts) and cellular processes (34,155 transcripts) were the most highly represented categories. Within molecular function subontology, binding (37,455 transcripts) and catalytic activity (32,678 transcripts) were amongst the most highly represented categories while within the cellular component subontology, the most transcripts are related to cells (35,816 transcripts) and organelles (35,769 transcripts).

The 97,454 transcripts were also annotated using the Kyoto Encyclopedia of Genes and Genomes (KEGG) database, which is a database of resources representing the functions and interactions of genes, and biological system at molecular level [19]. The KEGG database presents information as collections of manually drawn pathway maps [20]. Of the 97,454 transcripts, 18,158 were successfully mapped to the 153 KEGG pathways. Five main categories of KEGG pathways were successfully mapped, including metabolism, genetic information processing, environmental information processing, cellular processes and organismal systems. Of these five categories, metabolism was the most highly represented (with 82.2%), followed by organismal systems, genetic information processing, environmental information and cellular processes (S3 Fig).

### Identification of genes related to wood biosynthesis

In this study, only transcripts representing genes related to wood-formation biosynthesis were selected for SSRs and SNPs markers identification. Lignin biosynthesis-related genes consist of phenylalanine ammonia-lyase (PAL), cinnamate-4-hydroxylase (4CL), shikimate O-hydroxycinnamoyltransferase (HCT), caffeic acid O-methyltransferase (COMT), caffeoyl-CoA O-methyltransferase (CCoAOMT), cinnamoyl-CoA reductase (CCR) and cinnamyl alcohol dehydrogenase (CAD), while cellulose-biosynthesis-related genes consist of cellulose synthase (CesA), COBRA protein, KORRIGAN protein and α-L-fucosidase. Table 2 shows the number of transcripts related to genes involved in lignin and cellulose biosynthesis.

**Table 2.** Number of transcripts relation to genes involved in lignin and cellulose biosynthesis.

|  | **Enzyme**<br>**Symbol** | **Enzyme Commission**<br>**Number** | **Number of Transcripts** |
| --- | --- | --- | --- |
| **Lignin Biosynthesis** |  |  |  |
| Phenylalanine ammonia-lyase | PAL | [EC:4.3.1.24] | 16 |
| Cinnamate-4-hydroxylase | 4CL | [EC:1.14.14.91] | 17 |
| Shikimate O-hydroxycinnamoyltransferase | HCT | [EC:2.3.1.133] | 6 |
| Caffeic acid O-methyltransferase | COMT | [EC:2.1.1.68] | 13 |
| Caffeoyl-CoA O-methyltransferase | CCoAOMT | [EC:2.1.1.104] | 3 |
| Cinnamoyl-CoA reductase | CCR | [EC:1.2.1.44] | 17 |
| Cinnamyl alcohol dehydrogenase | CAD | [EC:1.1.1.195] | 9 |
| **Cellulose Biosynthesis** |  |  |  |
| Cellulose synthase | CesA | [EC:2.4.1.12] | 34 |
| COBRA protein |  |  | 3 |
| KORRIGAN protein |  |  | 12 |
| α-L-fucosidase | | [EC:3.2.1.63] | 14 |

### SNP marker discovery

In terms of putative single-nucleotide polymorphisms (SNPs) discovery, a total of 3,210,629 SNPs was detected from total dataset, and an average of one SNP per 32 bp was observed. Transition SNPs were found to be predominant (61.2%) as opposed to transversion SNPs (38.8%). Amongst transversion variations, G « T was the most highly represented (with 312,386 SNPs detected) followed by A « C (with 310,681 SNPs identified), C « G (with 310,376 SNPs identified) and A « T was the least common (with 309,289 SNPs identified) (Table 3).

**Table 3.** Overview of SNPs identified from three tissues and four rubber clones.

|  |  |
| --- | --- |
| Number of transcripts | 101,269 |
| Number of SNPs | 3,210,629 |
| SNP frequency | 1 per 32 bp |
| Transition | 1,967,897 |
| A « G | 983,962 |
| C « T | 983,935 |
| Transversion | 1,242,732 |
| A « C | 310,681 |
| A « T | 309,289 |
| C « G | 310,376 |
| G « T | 312,386 |

A more detail search was conducted to identify SNPs from genes related to cellulose and lignin biosynthesis. The lignin-biosynthesis pathway has seven gene toolboxes, while the cellulose-biosynthesis pathway has four gene toolboxes. A total of 1,786 SNPs was found from 11 lignin and cellulose gene toolboxes. Table 4 shows all the SNPs that were found in 11 gene toolboxes related to lignin and cellulose biosynthesis.

**Table 4.** Number of SNPs found in lignin- and cellulose-biosynthesis pathways.

| **Lignin Biosynthesis** | **Number of Single-Nucleotide Polymorphisms (SNPs)** |
| --- | --- |
| Phenylalanine ammonia-lyase (PAL) | 115 |
| Cinnamate-4-hydroxylase (4CL) | 227 |
| Shikimate O-hydroxycinnamoyltransferase (HCT) | 59 |
| Caffeic acid O-methyltransferase (COMT) | 108 |
| Caffeoyl-CoA O-methyltransferase (CCoAOMT) | 54 |
| Cinnamoyl-CoA reductase (CCR) | 152 |
| Cinnamyl alcohol dehydrogenase (CAD) | 84 |
| **Cellulose Biosynthesis** | **Number of Single-Nucleotide Polymorphisms (SNPs)** |
| Cellulose synthase (CesA) | 672 |
| COBRA protein | 16 |
| KORRIGAN protein | 149 |
| α-L-fucosidase | 150 |
| **TOTAL** | **1,786** |

### SSR Marker Discovery

The 101,269 transcripts were subjected to a search for putative SSR markers. A total of 14,956 SSRs were detected in 14,145 transcripts. A frequency of one SSR per 5.5 kb was identified. Dinucleotide SSRs have been reported to be the most abundant type of SSR in plant genomes. There were 12,767 di-, 2,172 tri-, 14 tetra- and 3 pentanucleotide potential SSRs. “AG/CT” motif was found to be the highest proportion, followed by “AT/AT” motif and “AAG/CTT” motif within rubber transcriptome dataset (S2 Table).

A detail search was conducted to discover SSRs from genes related to lignin and cellulose biosynthesis. A total of 31 SSRs have been found related to lignin and cellulose biosynthesis genes. However, after filtering the redundancy of SSRs within sequences covered by primers, the number of working primers was reduced to 18. The 18 SSR primers were selected from several genes including PAL, 4CL, COMT, CCR, CesA, COBRA and KOR (Table 5).

**Table 5.** SSRs markers and motifs identified from transcriptome of rubber clones related to lignin and cellulose biosynthesis genes.

| Genes | Number of SSR Primers | SSR motifs |
| --- | --- | --- |
| PAL | 1 | GT |
| 4CL | 3 | AG<br>AG<br>CT |
| HCT | 0 | - |
| COMT | 1 | CT |
| CCoAOMT | 0 | - |
| CCR | 1 | AC |
| CAD | 0 | - |
| CesA | 9 | AG<br>AG<br>AG<br>AG<br>AT<br>AT<br>ATT<br>GAT<br>GAT |
| COBRA | 1 | CT |
| KOR | 2 | GAT<br>GAT |
| α-L-fucosidase | 0 | - |
| **TOTAL** | **18** |  |

### SNP primers design and screening

SNPs found earlier from transcriptome datasets of genes related to lignin and cellulose biosynthesis were used to develop 764 primer pairs. However, after filtering the redundancy of SNPs within sequences covered by primers, the number of working primers was reduced to 151 primer pairs. All 151 SNP markers were amplified using conventional PCR with a set of rubber clones before they could be used as a marker for selection of a high-quality rubber clones. The validation process showed some positives results with approximately 68% primer pairs successfully amplified and amplicons were found producing expected band size on agarose gel. Of the 151 of SNP primers, 103 were successfully amplified SNP in the DNA sequence of rubber. Table 6 shows the total number of SNP primers generated and number of primers successfully amplified from genes related to lignin- and cellulose-biosynthesis pathways. A details list of SNPs primers can be found in S3 Table.

**Table 6.** Number of primers developed for SNP markers related to lignin- and cellulose-biosynthesis pathways.

| **Genes** | **Number of Primers** | **Number of Primers Successfully Amplified** |
| --- | --- | --- |
| PAL | 13 | 10 |
| 4CL | 26 | 13 |
| HCT | 5 | 4 |
| COMT | 12 | 12 |
| CCoAOMT | 7 | 2 |
| CCR | 18 | 10 |
| CAD | 6 | 4 |
| CesA | 55 | 48 |
| COBRA | 1 | 0 |
| KOR | 3 | 0 |
| α-L-fucosidase | 5 | 0 |
| **TOTAL** | **151** | **103** |

### SSR primer design and screening

The 18 SSR markers related to lignin and cellulose-biosynthesis gene toolbox were then selected for amplification using conventional PCR with a set of rubber clones before they could be used as a marker for selection of a high-quality rubber clone. All 18 SSR markers were successfully amplified and amplicons obtained were of expected size visualized on agarose gel. A details list of SSRs primers can be found in S4 Table.

### Phylogenetic analysis of genes related to lignin biosynthesis

In this study, phylogenetic analysis using DNA sequence can provide the evolutionary relationship between genes related to lignin biosynthesis among different rubber clones. Initially, the sequence alignment analysis was done to identify similarity regions of same genes from different taxa. The DNA sequence was initially compared to the online database available for sequence identification. The boundaries of COMT gene were determined by comparing with the sequences from other *Hevea* taxa obtained from GenBank (Accesion no.: JQ037840). The length of COMT gene for *Hevea* taxa varied from 884 to 895 bp. Moreover, the boundaries of CCoAOMT gene were determined by comparing with the sequences obtained from GenBank (Accesion no.: KU301753) where the length of CCoAOMT gene for *Hevea* taxa varied from 745 to 761 bp. Apart from that, the boundaries of CCR gene were determined by comparing with the sequences obtained from GenBank (Accesion no.: KU301753) where the length of CCR gene for *Hevea* taxa varied from 867 to 905 bp. In addition, the boundaries of CAD gene were determined by comparing with the sequences obtained from GenBank (Accesion no.: HQ229953). The length of CAD gene for *Hevea* taxa varied from 1,171 to 1,208 bp. The aligned sequence were then used as an input to construct phylogenetic tree.

We combined all the four gene sequences and subjected this new combined data to MP analysis. The maximum parsimony (MP) analysis of combined gene sequences has resulted in a data matrix of 3756 characters from 9 taxa including outgroup. From these 3756 characters, 2685 (71.5%) characters are constant while 597 (15.9%) characters are parsimony-uninformative and 474 (12.6%) characters are parsimony-informative. The 474 parsimony-informative characters produced tree length of 585 steps with a consistency index (CI) of 0.8900 and retention index (RI) of 0.8678. The resulting MP tree separated the four rubber clones into two clades (Fig 1). One clade consists of RRIM 2025 and RRIM 3001 while the other clade composed of PB 314 and RRIM 712. MP analysis has placed RRIM 2025 as a sister to RRIM 3001. With high bootstrap value of 96, the placement of RRIM 2025 next to RRIM 3001 was well supported. These two clones are latex-timber clones with larger trunk compared to PB 314 and RRIM 712 clones. Thus, the placement of RRIM 2025 next to RRIM 3001 are supported based on molecular and morphology characteristics. The other clade consists of PB 314 as sister to RRIM 712. MP tree has placed both clones next to each other based on the analysis of genes related to lignin biosynthesis with high bootstrap value of 89. Both clones also have relatively smaller trunk compared to RRIM 2025 and RRIM 3001.

**Fig 1.**
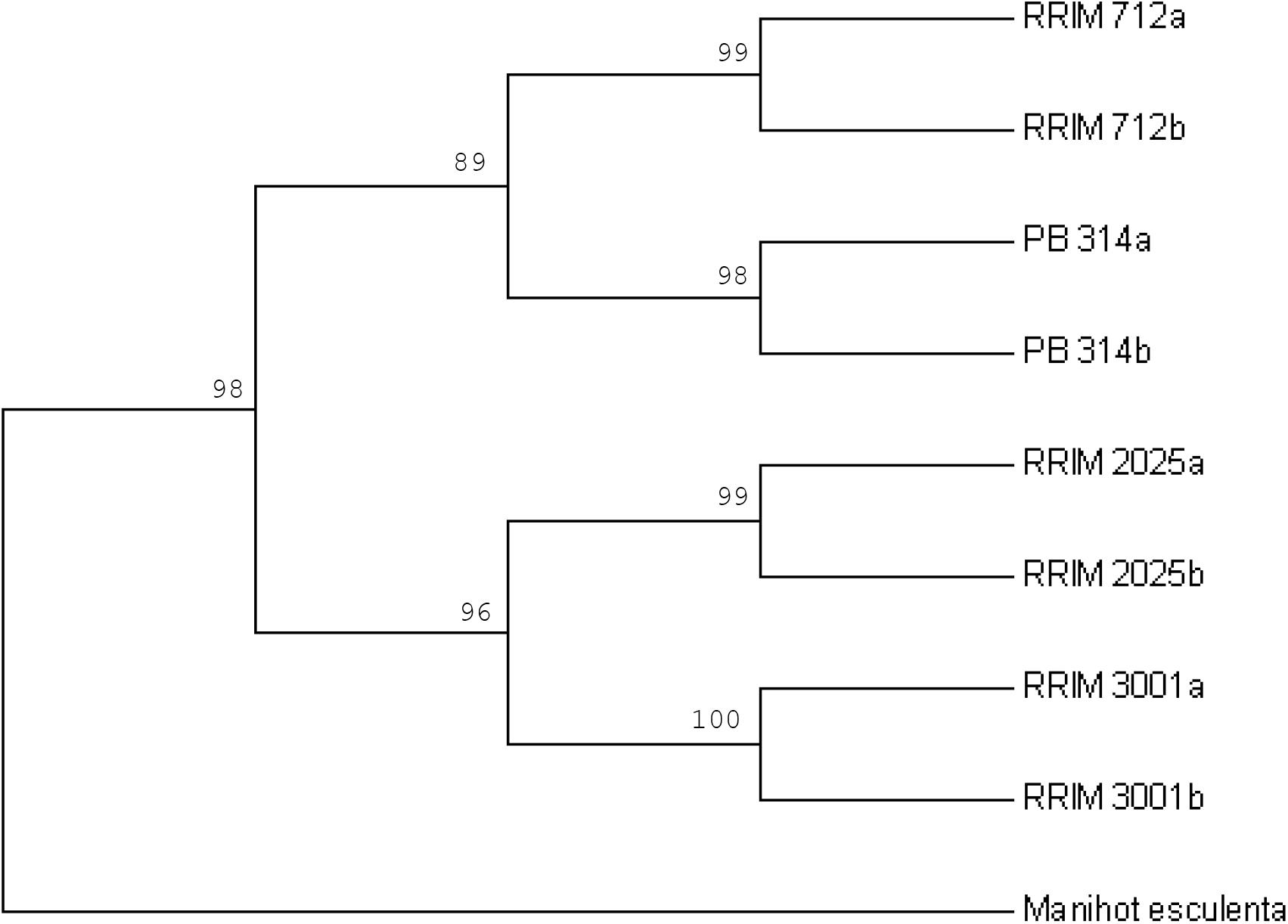
The maximum parsimony (MP) tree constructed from combines COMT, CCoAOMT, CCR and CAD genes. Values above branches are bootstrap values. COMT (caffeic acid O-methyltransferase); CCoAOMT (caffeoyl-CoA O-methyltransferase); CCR (cinnamoyl-CoA reductase); CAD (cinnamyl alcohol dehydrogenase).

## Discussion

Investigation of the genes related to rubberwood biosynthesis involves gathering information through transcriptome datasets at a molecular level. Understanding natural rubber production at a molecular level can bring new insights to programmes for rubber breeding. *H. brasiliensis* transcriptome data showed the mean length of transcripts to be 1,046 bp (N50 was 1,313 bp), and 101,269 transcripts were recovered in total. The transcriptome data can be accepted since similarities were found with previous studies in terms of *H. brasiliensis* transcriptome analysis. In our study, the mean length of transcripts was found to be far greater than that reported by [21], who used RRIM600 as their specimens and found the mean length of 19,708 transcripts to be 676 bp. In contrast, we found that N50 had 837 bp. Moreover, Mantello *et al*. [20] reported a higher number of transcripts (152,416), with lengths ranging from 97 bp to 13,266 bp within GT1 and PR255 clones. However, the mean length of transcripts (536 bp) and N50 were lower (720 bp) than our data. In addition, [22] reported that about 93.85% of their clean reads were successfully recovered, generating 98,697 transcripts with a mean length of 1,437 and 2,127 bp for N50. Recently, [23] reported that their study of CATAS93-114 generated 100,100 transcripts, with N50 having 2,048 bp.

Our annotation analysis indicated 83,771 hits in the NR database (83%), while 61,917 hits (61%) were found in the UniProtKB/Swiss-Prot database. Our annotation analysis showed higher percentages than those of various other studies, as follows: [20]: 63% (NR database) and 47% (UniProtKB/Swiss-Prot database; [22]: 65% (NR database) and 46.9% (UniProtKB/Swiss-Prot database); [24]: 72.6% (NR database) and 55.2% (UniProtKB/Swiss-Prot database; and [25]: 66.3% (NR database) and 42.1% (UniProtKB/Swiss-Prot database). Moreover, annotation analysis using the GO database found that amongst the three main subontologies, biological processes were the most highly represented, followed by molecular functions and cellular components. Our results are also similar to those described by [21] and [26]. However, [24] suggested that cellular components were the most highly represented, followed by biological processes and molecular functions, while [25] reported that biological processes were the most highly represented ontology followed by cellular components and molecular functions. Furthermore, annotation analysis using the KEGG database showed that about 18% of the transcriptome dataset was successfully mapped onto 153 pathways. Most of the transcripts were found to be involved in metabolic processes, followed by organismal systems, genetic information processing, environmental information, and cellular processes. The percentage mapped using the KEGG database was similar to that described by [20], with 17.1% (137 KEGG pathways), and [22], with 15.4%. However, [24] and [25] reported that the total transcriptome dataset mapped to the KEGG database was slightly higher than our data, with 53.1% (123 pathways) and 39.9% (128 pathways) being successfully mapped.

A total of 3,210,629 SNPs and 14,956 SSRs associated with potentially important genes were identified from the transcriptome dataset. The high number of SNPs from transcriptome data indicated the high allelic variability present in rubber trees that is able to be detected using this NGS technology. Moreover, about 73% from all SNP markers developed from lignin and cellulose biosynthesis pathway were successfully amplified on the random rubber clones. Our results have a higher density than those described by [20] (1 SNP per 125 bp), [27] (1 SNP per 178 bp) and [28] (1 SNP per 1.5 kb) for the rubber trees studied. Apart from that, the high percentage of working primers showed that SNPs are present in various rubber clones and may be involved in rubber tree physiologies. Furthermore, the 18 SSR primers which were successfully generated and amplified could differentiate and showed a high frequency of polymorphisms between rubber clones. Although SNP markers are the most abundant type of DNA polymorphism in genomic sequences, SSR together with SNP could be the best candidate markers for genetic breeding and to investigate the variability of genes related to rubberwood biosynthesis in rubber trees.

Usage of biomarkers in breeding programmes could help increase the chances of producing new high-yield rubber clones without a high cost and in a shorter time. These primers can be used as a marker to get information about whether new rubber clones have potentially high wood volumes and can be grown and supplied to rubberwood industries. These markers can also be used by rubber breeders to obtain information about newly bred trees as young as one or two years old, and this could play a vital role in rubber breeding programmes.

## Conclusions

The use of biomarkers in rubber breeding programmes (including using single-nucleotide polymorphisms (SNPs) or simple-sequence repeats (SSRs) could help rubber breeders determine the characteristics of new rubber clones in less time, without having to wait 30 years for a clone to be released. Using conventional polymerase chain reaction (PCR) in the validation process could help achieve targets faster than ever. Successfully developed 103 SNPs and 18 SSR primer sets could be used in genetic diversity studies and rubber breeding programmes focusing on rubberwood formation. Future studies could, therefore, provide an insight into how rubberwood quality and production processes could be improved.

## Acknowledgements

This project is funded by RISDA (304/PBIOLOGI/650728/P137) awarded to Ahmad Sofiman Othman of Universiti Sains Malaysia. We thank Yue Keong Choon (Universiti Sains Malaysia) for collecting the samples used in this work and Mohd Khairul Luqman Mohd Sakaf (Universiti Sains Malaysia) for assisting in analysis.

## Supporting information

**S1 Table. Raw reads statistics of transcriptome analysis in rubber clones**.

**S2 Table. SNPs loci identified from transcriptome of bark, leaf and latex tissues from RRIM 3001 related to lignin biosynthesis**.

**S3 Table. SNP primers for lignin and cellulose-biosynthesis related genes**.

**S4 Table. SSR primers for lignin and cellulose-biosynthesis related genes**.

**S1 Fig. Top hits species distributions against NR database using BLASTx program. *Manihot esculenta* is the top hits species followed by *Jatropha curcas, Ricinus communis, Hevea brasiliensis* and *Populus trichocarpa***.

**S2 Fig. Summary of GO classification of *H. brasiliensis*. Biological process is the most highly represented category followed by molecular functions and cellular components**.

**S3 Fig. Summary of transcriptome annotations against KEGG database. Metabolism is the most highly represented followed by organismal systems, genetic information processing, environmental information, and cellular processes**.

